# Sex and pedagogy influences in physics learning-related reorganization of brain activation

**DOI:** 10.1101/791301

**Authors:** Jessica E. Bartley, Michael C. Riedel, Taylor Salo, Katherine L. Bottenhorn, Emily R. Boeving, Robert W. Laird, Matthew T. Sutherland, Shannon M. Pruden, Eric Brewe, Angela R. Laird

## Abstract

Physics is a challenging academic pursuit in which university students regularly struggle to achieve success. Female students tend to perform negatively on introductory physics conceptual assessments compared to their male peers; however, active-learning classroom curricula are known to broadly improve performance on these tests. Here, we used fMRI to delineate physics-related brain activity in 107 students and probed for changes following a semester of active-learning or lecture-based physics instruction. Large-scale reorganization of brain activity accompanying learning occurred in a mixed frontoparietal and default mode network. Sex differences were observed in frontoparietal, default mode, and primary visual areas before and after instruction. Regions showing significant pedagogy, sex, and time interactions were revealed during physics retrieval, suggesting the type of class students complete may influence sex differences in how students retrieve information. These results reveal potentially elucidating sex and pedagogy differences underlying the neural mechanisms supporting physics learning.

## INTRODUCTION

University education plays a critical role in cultivating and training the next generation of science, technology, engineering, and mathematics (STEM) professionals. Yet many students from underrepresented groups experience challenges that pose barriers to their education^1^, a problem that contributes to a relative disproportion of women in STEM^2^. This issue is particularly evident in physics, with only 21% of baccalaureate degrees awarded to women in the United States^3–6^. Female students tend to underperform on physics conceptual assessments relative to their male peers^7^, which has been linked to a range of social factors (e.g., attitudes and beliefs about physics, experiences of stereotype threat, differences in preparation and exposure to physics concepts, career expectations, science identity, and perceptions of belongingness in physics classrooms^7–10^). While this prior work suggests evidence of sex differences associated with students’ experiences in introductory physics, it is unknown whether such differences may translate to distinctions in the underlying neurobiology that supports physics learning.

Functional neuroimaging investigations assessing how individuals conceptualize, reason with, and learn about physical systems indicate physics cognition engages multiple brain areas across a fronto-parietal network (FPN)^11–13^ similar to that of domain-general problem solving^14^. Additionally, classroom learning interventions have been shown to increase functional connectivity in task-critical reasoning- and retrieval-related brain systems following skill acquisition^15,16^, and evidence indicates enhanced cross-system autonomy accompanying learning^17^. However, no study to date has examined putative neurobiological changes associated with physics classroom learning. It is also currently unknown how different aspects of physics cognition, including reasoning and content knowledge retrieval, may differently engage the FPN and associated networks. Furthermore, it is unclear the degree to which observed sex differences in physics performance across these measures are accompanied by differences in brain function.

Given the impact of sociological differences in classroom experience on conceptual performance, any meaningful characterization of learning-related brain function ought to consider not only the content learned but also how different environments equitably support that learning. If physics instruction does result in a reorganization of brain activity, then different pedagogies – especially those that build supportive learning communities as part of their curriculum – may affect these changes in sex-specific ways. Active-engagement curricula (i.e., active-learning instruction that directly involves all students in the learning process) improve conceptual test scores and odds of success relative to lecture-based instruction for all students^18,19^. They also support the positive development of female students’ physics self-concepts, science identities, and attitudes and beliefs about physics^20–24^, suggesting active-learning may be both broadly effective as well as specifically beneficial for female students’ success. Physics Modeling Instruction (MI) is one such active-learning pedagogy wherein students engage with instructors and peers in studio classrooms to develop, test, and verify physics models through experimentation and collaborative inquiry-based group activities^25^. While the benefits of MI relative to traditional lecture instruction (LI) have been well documented^21,26–29^, any accompanying neural differences are unknown. Understanding how these academic environments might differentially support physics learning-related brain function may shed light on potential sex- and pedagogy-related influences in physics learning beyond the interpretations available with behavioral and cognitive studies alone.

Here, we sought to characterize the potential influences of sex and pedagogy on physics-related brain activity, determine how classroom learning is associated with functional reorganization of large-scale brain networks in physics students, and investigate if pedagogical approach differentially impacts these shifts in female and male students. We used functional magnetic resonance imaging (fMRI) to delineate physics reasoning- and retrieval-related brain networks in 107 university-level introductory physics students (48 female and 59 male students) and probed for differences resulting from a semester of active learning and lecture-based physics instruction. Students underwent MRI scanning before and after a 15-week long MI or LI physics course and completed in-scanner tasks based on the Force Concept Inventory^30^ (FCI; measuring physics conceptual reasoning), a physics content knowledge test (PK; probing retrieval of physics classroom content), and a content-general transitive inference control task (TI; examining general reasoning to assess the domain specificity of physics instruction-related shifts in brain activity) (**Figure 1**). Across all students, we anticipated increased activity in the fronto-parietal network at post-instruction compared to pre-instruction for the physics-specific FCI and PK tasks but expected no shifts in the domain-general TI task. Given the pedagogical differences associated with Modeling Instruction compared to traditional Lecture Instruction, we further hypothesized that MI would result in increased learning-related activity associated with reasoning and critical thinking, whereas LI would be associated with retrieval of facts and formulas stored in semantic memory. Lastly, while we hypothesized that these classroom differences would be broadly observed for both female and male students, but we predicted an interaction between sex and pedagogy yielding sex-specific patterns of brain reorganization. Our overall objective was to shed new light on how neural changes enable learning in a fundamental STEM discipline and to identify and guide action supporting learning that is inclusive for all students.

**Figure 1.**
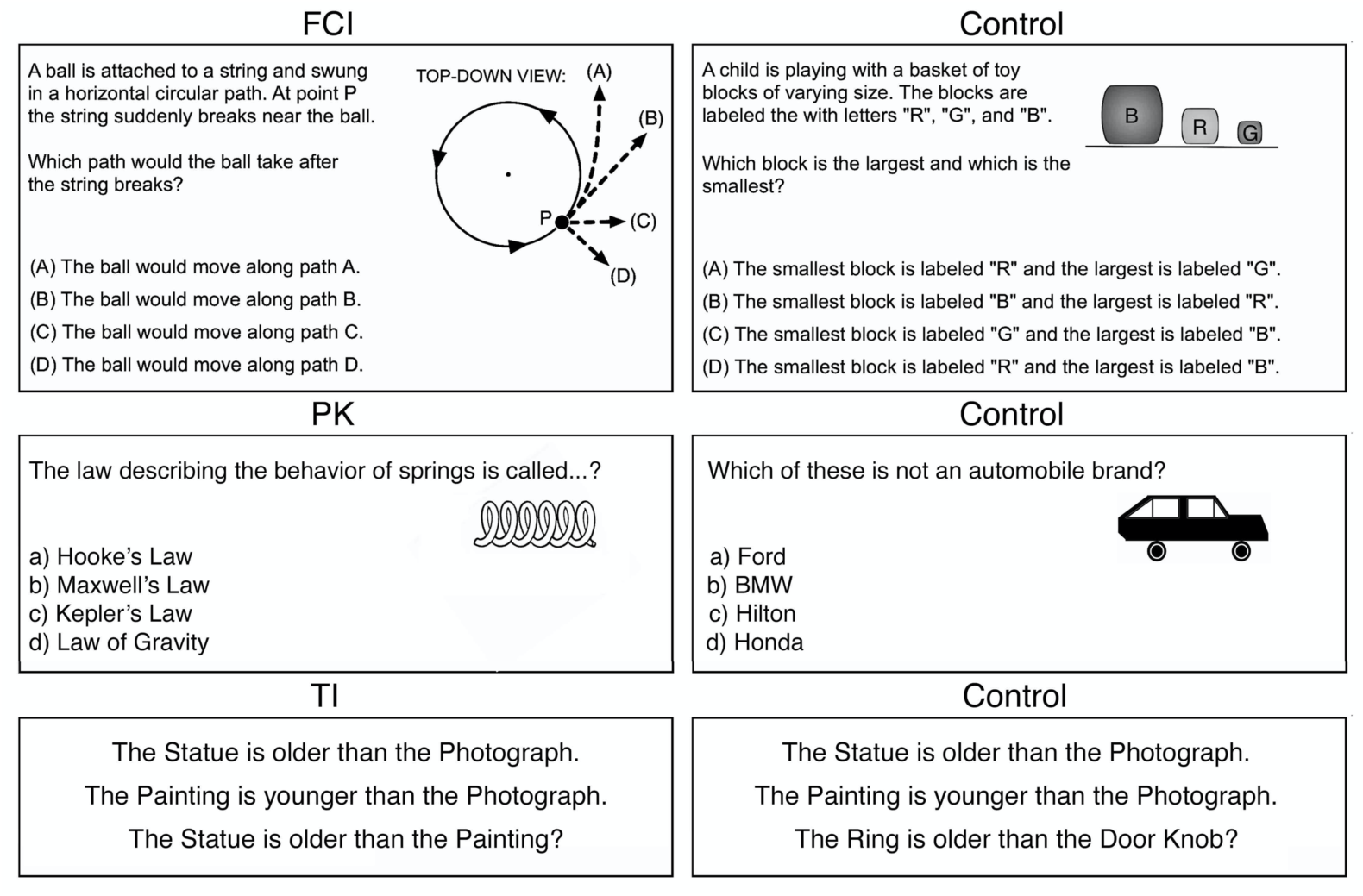
Study Design and MRI Tasks. Representative questions from each of the three in-scanner tasks: the Force Concept Inventory (FCI) contrasted a physics reasoning condition with a high-level perceptual control condition, the physics knowledge (PK) task contrasted a physics knowledge condition with a general knowledge condition, and the Transitive inference (TI) task contrasted a general reasoning condition with a baseline control condition.

## RESULTS

### Behavioral measures indicative of physics learning

Students scored significantly better at the post-instruction stage on both FCI and PK stimulus questions, but not on TI questions (FCI: Pre = 46%, SE_Pre_ = 2.1%, Post = 61%, SE_Post_ = 1.9%, *P* < 0.0001; PK: Pre = 66%. SE_Pre_ = 1.2%, Post = 77%, SE_Post_ = 1.2%, *P* < 0.0001; TI: Pre = 83%, SE_Pre_ = 1.2%, Post = 85%, SE_Post_ = 1.0%, *P* = 0. 053), indicating students successfully learned to reason and recall facts about physics concepts across instruction. No major shifts in the content-general TI condition (on which students received no instruction across the course) were expected. A mixed effects analysis of variance (ANOVA; Instructional Group (LI, MI) x Sex (Male, Female) x Time (Pre, Post)) of task accuracy yielded significant main effects for both sex and time (FCI Accuracy: *F*_*sex*_(1, 104) = 32.2, *P* < 0.0001, Cohen’s d = 0.89; *F*_*time*_(1, 104) = 66.1, *P* < 0.0001, d = 1.02; PK Accuracy: *F*_*sex*_(1, 104) = 25.7, *P* < 0.0001, d = 0.41; *F*_*time*_(1, 104) = 74.7, *P* < 0.0001, d = 0.73) but not for class (FCI Accuracy: *F*_*class*_(1, 104) = 2.24, *P* = 0.138, d = 0.43; PK Accuracy: *F*_*class*_(1, 104) = 4.86, *P* = 0.0297, d = 0.37) (**Figure 2**). No significant interactions in accuracy were observed between class, sex, and time. *Post hoc* Tukey tests confirmed FCI and PK scores significantly differed between pre- and post-instructional sessions as well as between female and male student groups at *P* < 0.0001. Male students scored higher than female students at both time points (males: FCI Pre = 55%, FCI Post = 68%, PK Pre = 72%, PK Post = 80%; females: FCI Pre = 36%, FCI Post = 52%, PK Pre = 62%, PK Post = 72%), indicating students entered the physics courses with an existing achievement gap and, while both female and male students successfully learned physics content, the course failed to reduce the initial performance difference. On average, students received a final passing grade of 2.87 out of 4.00 (B-range; see SI for grading scale details; **Supplementary Figure 1**) across all physics courses. There was no significant difference in physics course grade between female and male students (*M*_*male*_ = 3.01, *SD*_*male*_ = 0.98, *M*_*female*_ = 2.69, *SD*_*female*_ = 1.06; *P* = 0.118), matching previous observations^31^, which may be due to grades consisting of multiple performance measures (e.g., exams, homework, attendance). There was a small but significant difference between the course grades of LI and MI students, with MI students achieving higher grades than their LI peers (*M*_*LI*_ = 2.66, *SD*_*LI*_ = 1.14, *M*_*MI*_ = 3.06, *SD*_*MI*_ = 0.87, *P* = 0.048), consistent with previous findings^18^. Collectively, these results indicate students successfully learned physics content across instruction, yet sex-specific effects may have influenced this learning process and different course types may have differentially supported student’s academic success.

**Figure 2.**
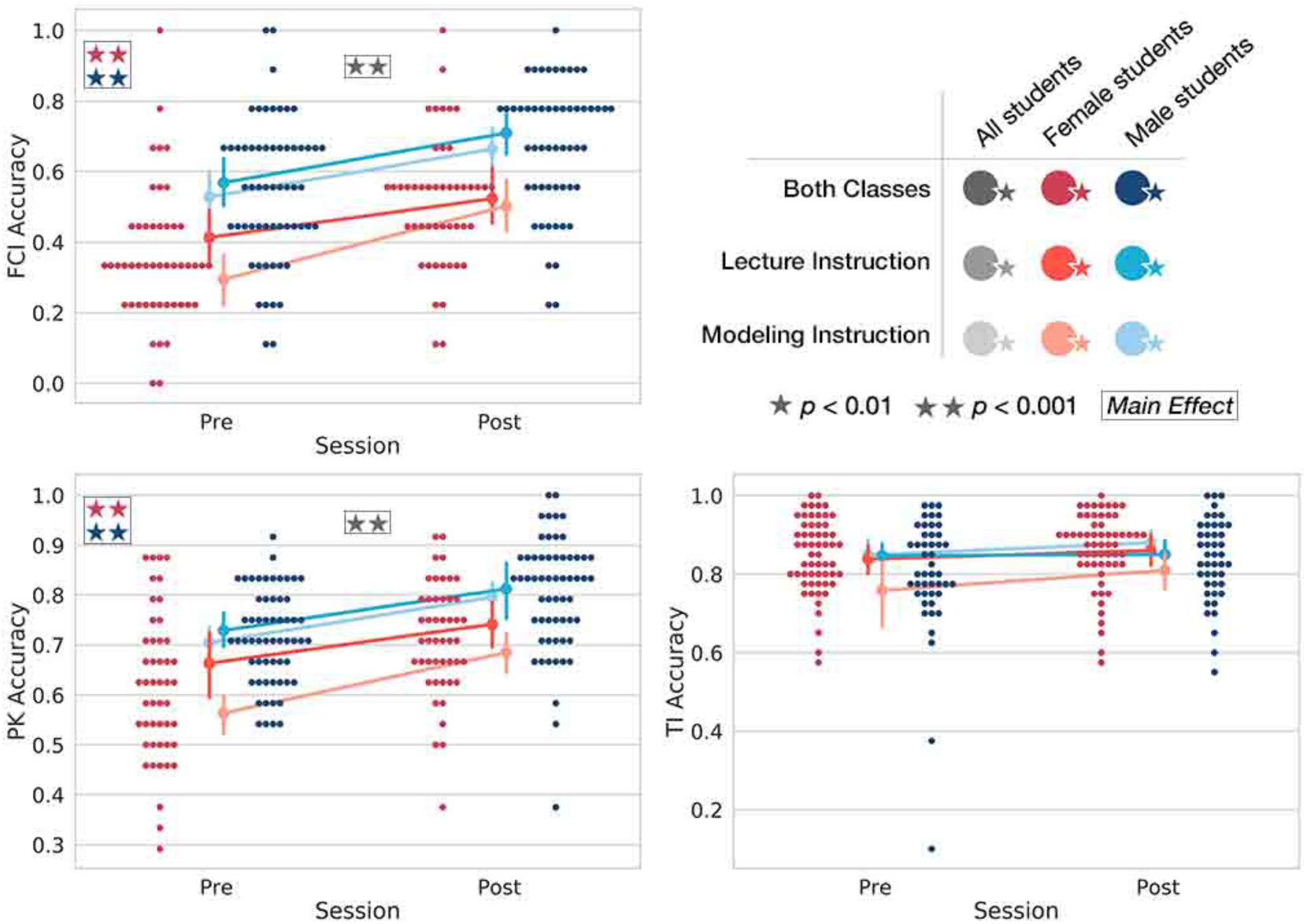
In-Scanner Task Accuracy. Scatter plots show pre- to post-instruction shifts in task accuracy for Female and Male students for the FCI (top left), PK (bottom left), and TI (bottom right) tasks. Trend lines depict the shifting mean of each class and sex group. Mixed effects ANOVA (Instructional Group x Sex x Time) of task accuracy yielded significant main effects for both sex and time in the FCI and PK tasks, as indicated by red/blue and gray stars, respectively. No significant time or sex effects were observed for the domain-general TI task.

### Learning-related differences in physics brain activity

Pre- and post-instruction brain activity was observed across multiple FPN variants for the FCI and PK physics tasks as well as for the TI general reasoning task (**Supplementary Figure 2; Supplementary Table 1**). We found learning-related changes in brain activity across all students within a mixed FPN and default mode network (DMN) associated with both physics tasks, but not with the TI task (**Figure 3**; **Supplementary Figure 3 and Table 3**). Significant increases in FCI- and PK-related brain activity following instruction were observed across the prefrontal cortices (PFC) with clusters along the inferior precentral sulcus, in the dorsomedial PFC (dmPFC), dorsolateral PFC (dlPFC), bilaterally in the frontal poles, and in the ventromedial PFC (vmPFC). Parietal and posterior medial areas also showed increased physics-related activation after instruction in the posterior parietal cortices (PCC), retrosplenial cortex (RSC), precuneus, as well as in bilateral and left-emphasized horizontal intraparietal sulcus (IPS) and angular gyri. Additionally, the PK task showed significant increased activity post-relative to pre-instruction in the caudal portion of the left lateral occipitotemporal gyrus. These findings reveal notable and pronounced agreement in increased large-scale brain activity across both independent physics tasks following classroom instruction. Importantly, no significantly increased activity was observed during the domain-general TI task, indicating physics-related increased FPN-DMN brain function subtends physics content learning as resulting from classroom instruction. Additionally, a whole-brain parametric modulation analysis of learning-related increases in brain activity with course grade yielded no significant results, indicating the relationship between physics course grade and task-related increases in brain activity may not be directly proportional. Additional explanatory factors may be necessary to more fully model the link between success in physics course grade and learning-related brain function.

**Figure 3.**
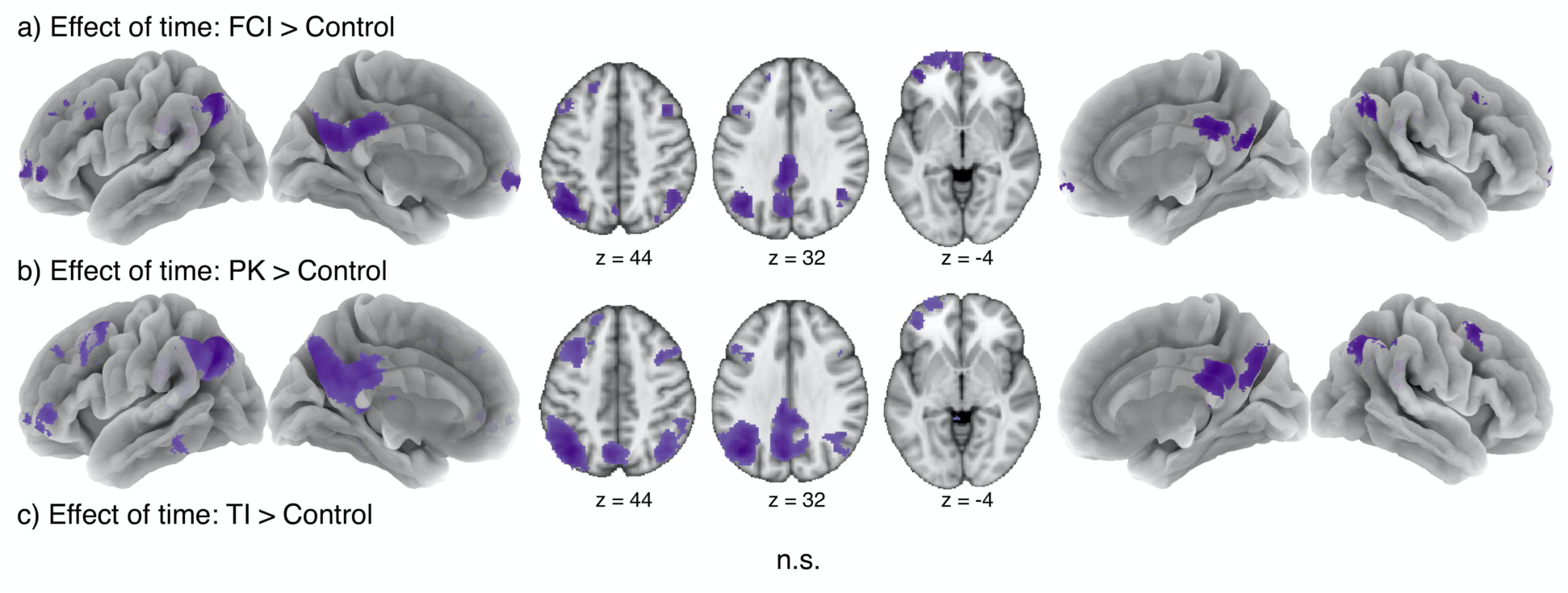
Changes in Activation Across Instruction. a) Post > Pre: FCI > Control, b) Post > Pre: PK > Control, and c) Post > Pre: TI > Control (i.e., no significant effects were observed for the domain-general task). Activation maps were thresholded using a cluster defining threshold of P < 0.001 and a cluster extent threshold of P < 0.05, FWE corrected.

### Sex influences in learning-related brain activity

To investigate sex influences in physics brain activity we probed for group similarities and differences in FCI- and PK-related task activity at both time points. Both female and male student groups activated highly similar pre- and post-instruction FPNs during the FCI and PK tasks (**Figure 4a**). However, significantly greater physics-related brain activity was also observed for male students at both time points in areas across an FPN similar to that of the overall FCI and PK networks, and for female students in DMN and visual-related areas (**Figure 4b**). Male students showed significantly greater FCI-related activity in frontal, parietal, and occipitotemporal regions (dlPFC, lateral orbitofrontal cortices (OFC), horizontal IPS, inferior parietal lobule, V5/MT+; areas similar to those revealed by the overall FCI > Control contrast) compared to female students. Female students more significantly activated visual and medial posterior areas (PCC, precuneus, cuneus, lingual gyri) at both time points during the FCI task, as well as increased frontal (medial frontal gyrus, right precentral gyrus, right inferior frontal gyrus) and anterior insular activity before instruction, compared to their male peers. These effects did not significantly change across instruction and did not differ across class types. Reduced but similar sex differences were observed during the PK task at pre- and post-instruction stages. Extensive primary visual cortex PK-related activity was observed in female vs. male students, and left FPN (dlPFC, horizontal IPS, inferior parietal lobule) PK-related activity occurred in male vs. female students at both time points. Additionally, increased PK-related activity at post-instruction occurred in bilateral temporal poles for female vs. male students and increased pre-instruction PK-related activity was detected in the right occipital cortex, bilateral fusiform gyri, and cerebellum for male vs. female students.

**Figure 4.**
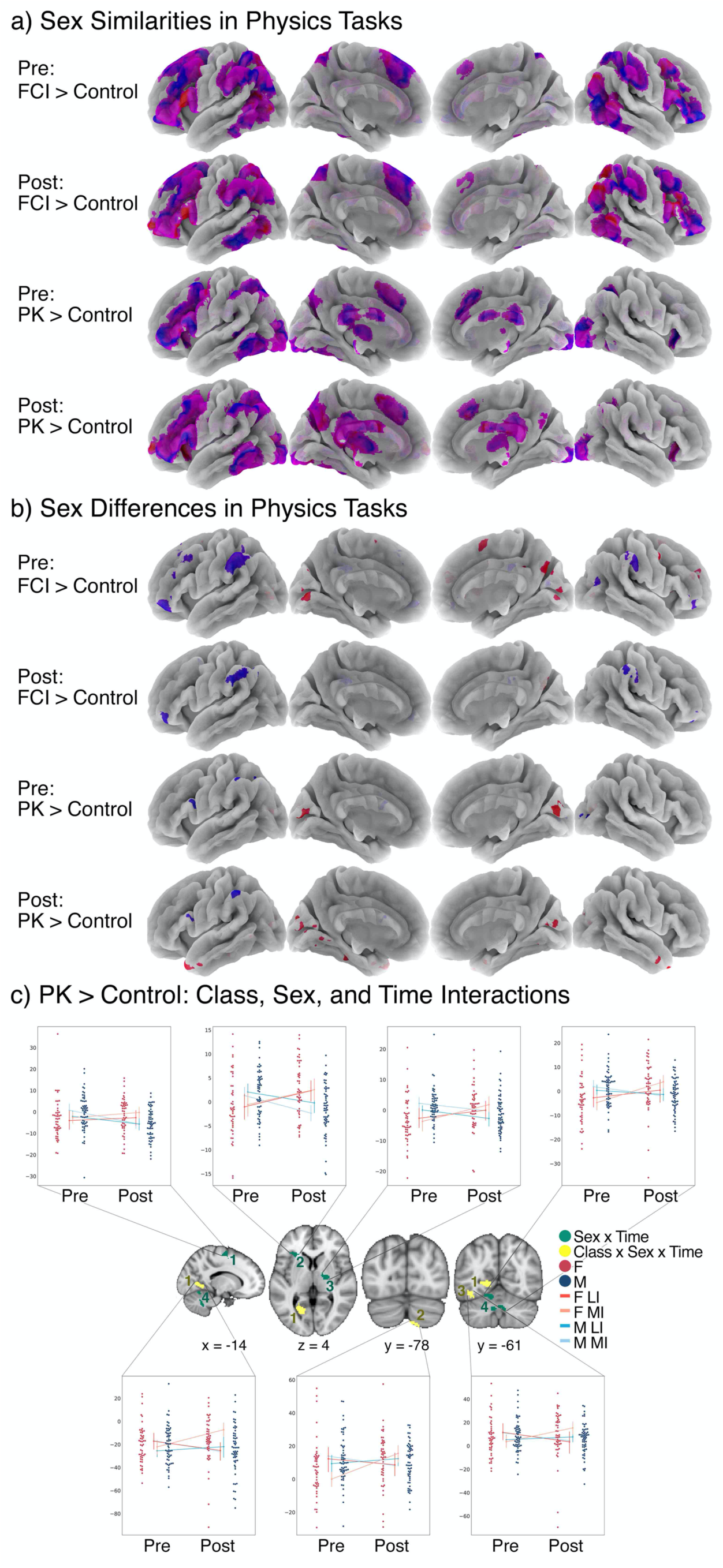
Sex and Pedagogy Influences in Physics Learning-Related Shifts of Brain Activation. a) Sex similarities in FCI > Control and PK > Control at pre- and post-instruction stages; group activity for female students is depicted in red, male students in blue, and regional overlap between female and male group activity in purple. b) Sex differences for Female > Male (red) and Male > Female (blue) contrasts for each physics task is shown for pre- and post-instruction stages. c) Regions exhibiting significant two-way (sex x time) interactions (green) and three-way (pedagogy x sex x time) interactions (yellow) are shown for PK > Control. Associated swarm plots of subject-level PK > Control beta values within these regions indicate the direction of each interaction. Activation maps were thresholded using a cluster defining threshold of P < 0.001 and a cluster extent threshold of P < 0.05, FWE corrected.

### Pedagogical influences in learning-related brain activity

Next, to investigate pedagogical influences in physics brain activity, we probed for group similarities and differences in FCI- and PK-related brain activity at post-instruction. No significant differences were observed across MI and LI groups for the FCI > Control and PK > Control contrasts after physics instruction.

### Interactions between sex, pedagogy, and time

Finally, to test for significant group differences in how physics learning-related brain activity shifted across time, potentially in sex-specific ways, we performed a group-level whole-brain three-way ANOVA (Instructional Group (LI, MI) x Sex (M, F) x Time (Pre, Post)) to probe for potential interactions between pedagogy, sex, and/or time. Two-way PK task interactions occurred between sex and time in the cerebellum, thalamus, anterior insular cortex, and premotor cortex and three-way PK task interactions occurred between pedagogy, sex, and time in the cerebellum, fusiform gyri, and lingual gyri. Region of interest (ROI) inspection indicated PK task activity increased in female students but decreased in male students across instruction in all two-way interaction clusters (**Figure 4c, green**). Critically, female MI students also showed considerable increases in PK-related activity across instruction in all three-way interaction clusters (**Figure 4c, yellow**). In these same ROIs male LI students showed smaller increases in activity, while both male MI and female LI students exhibited decreases in activity post-relative to pre-instruction. These findings indicate female and male students may engage brain areas differently during physics thinking, and these differences may depend on whether they completed a traditional or active learning physics class.

## DISCUSSION

Our results show pre- and post-instruction physics-related cognition engages similar FPN activity in both female and male students. Moreover, we observed common neurobiological changes in all students in a mixed FPN-DMN system across physics classroom instruction. These shifts, accompanied by improved task accuracy scores, suggest FPN-DMN coupling may play a critical role in physics learning. Given known instruction-related performance differences and the sex achievement gap, we had anticipated main effects for both sex and class type. In support of the first hypothesis, we detected sex differences in physics-related brain activity at both time points, with increased DMN and visual activity in female students and FPN activity in male students. These results indicate that while sex similarities supporting physics-related brain function are prominent, specific regions may also be differently engaged in sex-specific ways. No main effects of pedagogy in brain activity were detected, contrary to our secondary hypothesis. Instead, we observed significant sex x time and pedagogy x sex x time interactions associated with physics content knowledge retrieval. These results shed light on a potential and previously unmeasured neural basis for differences coincident with the existing achievement gap that classroom instruction regularly fails to reshape. Our findings suggest that sex differences in physics learning-related brain activity are indeed malleable and, critically, may shift differently across instruction depending on whether students completed a traditional or active learning physics class.

### Classroom learning is accompanied by large-scale reorganization of physics-related brain activity

Consonant longitudinal increases in a mixed FPN-DMN brain system occurred across classroom learning in both physics tasks. Comparatively, no such increases were detected during general reasoning. Given the backdrop of no TI effects, together with improved physics scores, our findings indicate FPN-DMN integration may be critical for successful, learning-informed physics cognition. Interestingly, cognitively demanding FPN-supported tasks are frequently known to suppress DMN activity^32^, yet our results suggest physics cognition after instruction evokes cooperation between these two systems. These findings are consisted with the Arousal, Balance, and Breadth of Attention model for whole-brain dynamics^33^ in which PCC engages multiple brain systems in time-dependent interactions during tasks where the attentional focus is sufficiently broad. According to this model, shifting connectivity between the PCC and various whole-brain systems engenders adaptive monitoring and responses to behaviorally relevant stimuli occurring outside the cognitive task, thereby rapidly changing the cognitive state according to the task’s needs^33^. In our sister paper focused on the FCI task^12^, we examined post-instruction physics-related brain dynamics and found joint FPC-DMN activity supported decision making, perhaps indicating physics thinking relies on mental exploration and episodic memory retrieval during solution generation^12,34,35^. Our current findings indicate these reasoning-related processes may also generalize to those that support physics content knowledge retrieval across learning.

### Sex and pedagogy influence physics learning-related brain activity

Students entered the physics courses with an existing achievement gap that favored males, consistent with previous findings^7^. Physics instruction (both LI and MI) failed to eliminate this gap, and these disparities were paralleled by sex differences in physics-related brain activity, with male students showing increased activity in FPN areas and female students showing increased activity in DMN and visual areas. We note that these findings do not support a gender essentialist interpretation of sex-specific dissociations in physics-related brain activity in which disparities are attributed to inherent biological differences. Indeed, considerable sex similarities broadly across the FPN indicate the ways in which female and male students similarly engage in physics-related cognition. Given the various factors thought to influence the performance gap (e.g., differences in preparation or prior exposure to STEM content^7,36^, gendered differences in science identity^9^, self-efficacy^37^, stereotype threat^7,38^, and implicit bias^7,36^), it is more likely that neurobiological distinctions arise as extensions of, or interactions with, multiple external environmental and socioemotional factors. Additional research is needed to better connect findings in neuroscience with educational and socioemotional research on learning.

Importantly, while the achievement gap persisted across instruction, we nonetheless detected regions displaying significant interactions between sex and time, as well as pedagogy, sex, and time during physics retrieval, suggesting sex-related differences in physics cognition can indeed shift as the result of thoughtful pedagogy. This establishes valuable insight into how physics instruction can impact students’ brain function in sex-specific ways, and thereby how instructors may be able to use pedagogy to target existing disparities. Three-way interaction clusters were located in visual areas of the brain, suggesting differences in how visualization of physics concepts are taught may differently influence students learning-related brain function. Female modeling students showed significantly increased activity in these visual clusters relative to all other groups. This may be due to MI’s increased emphasis on the use of multiple visual representations of physics concepts. Under this interpretation, it is possible that the sex differences in learning-related brain activity may be a manifestation of classroom differences. We note that our findings do not indicate any one set of neural support is more effective than another, but we do observe that how physics students learn to visualize appears to be of particular importance across instruction. Students may be learning these skills differently based on what pedagogical approach they have undergone, combined with their different gendered experiences during learning. Physics instructional emphasis may benefit student by more targeted instruction, especially increased emphasis of visualization techniques, in order to impact sex-related differences.

### Limitations

We acknowledge that our results may be specific to conceptual physics thinking and may not generalize to mathematically rigorous forms of physics problem solving, but we note that conceptual physics cognition likely forms the basis for more mathematically complex forms of physics reasoning. We also note that the present study focused on sex, but not gender (i.e., how students identify as women, men, or non-binary) differences. Given the wealth of environmental differences women and men experience across development, gender may be a more appropriate target for assessing dissociations in physics thinking than biological sex. Future studies may benefit by considering how students differently gendered behaviors and/or experiences may impact the neural mechanisms of physics learning. In particular, measures of academic motivation^39^, STEM anxiety^40–43^, attitudes and beliefs about science^44–46^, and environmental experiences may help delineate the role of behavior and environment have in the interplay between brain regions supporting physics learning and academic performance.

### Impact on instruction

We found that consistent patterns of brain activity underlying student physics thinking indicate executive function, visualization, and default mode processing are of critical importance to both female and male students. We additionally found that classroom instruction yields large-scale shifts in physics-related activity, but preexisting differences in how female and male students engaged FPN, DMN, and visual areas during physics thinking were left largely unaffected across instruction. Multiple brain regions exhibited sex and pedagogy-specific shifts in physics content knowledge-related activity, thus indicating existing sex differences in physics thinking are pliable through thoughtful pedagogical interventions. Importantly, while behavioral research has revealed inequities in success and experience, these observations have yet to provide insight on how to effectively guide action. Here, we propose a new emphasis on instructional visualizations in physics, especially towards teaching visualization techniques during physics cognition and problem solving. This may be accomplished, in part, by emphasizing the initial drawing of the problem or making increased use of in-class demonstrations. We note that, given the substantial sex similarities observed, our results do not support an interpretation in favor of sex-segregated classes. Rather, physics teachers may benefit their students by considering student’s differently gendered experiences in their classes and consider how innovative pedagogies may be used to support their success.

In conclusion, the current study strives to elucidate the relationship between physics learning and physics-related changes in brain activity. By grounding neuroscience studies of learning in educational theory and pedagogy, instructors and researchers can begin to edify the extent to which neurobiological changes are supported by intrapersonal and environmental factors. We can thus work to clarify, define, and create new models of learning that provide insight into the underpinnings of learning difficulties and how to prevent them^47–49^.

## METHODS

### Participants and study design

One hundred and twenty-two healthy, right-handed undergraduate students took part in this study. One hundred and seven of these participants completed both pre- and post-instruction MRI scans. This included 48 female students (range = 18-25 years, M = 19.87 years, SD = 3.26 years) and 59 male students (range = 18-25 years, M = 20.09 years, SD = 1.46 years). Participants were first-time enrollees in a semester of college-level, calculus-based introductory physics at Florida International University in Miami, Florida. Course content emphasized problem solving skill development and covered topics in classical Newtonian mechanics, including motion along straight lines and in two and three dimensions, Newton’s laws of motion, work and energy, momentum and collisions, and rotational dynamics. A total of 51 students were enrolled in traditional Lecture Instruction (LI) physics classes in which a professor delivered physics lectures for the majority of class time. The remaining 56 students completed an active-learning Modeling Instruction (MI) physics class set in a studio-style classroom. The MI course involved increased experimentation with physical systems and engagement with instructors and peers wherein students completed collaborative, inquiry-based activities to develop, test, and verify models about physical phenomena.

All participants self-reported to be free from cognitive impairments, neurological and psychiatric conditions, and use of psychotropic medications. Students completed one beginning-of-semester (“pre-instruction”) fMRI session and a second identical end-of-semester session (“post-instruction”) following the conclusion of the 15-week semester. Pre-instruction data were acquired no later than the fourth week of classes and before the first course exam, and post-instruction sessions were completed no more than two weeks after the physics final exam. All 107 participants (56 MI; 48 female) completed the FCI and PK tasks at both time points and 103 participants (53 MI; 45 female) completed the TI task at both time points. Written informed consent was obtained in accordance with FIU Institutional Review Board approval.

### MRI tasks

Participants completed the Force Concept Inventory (FCI^50^) physics reasoning task^12^, a physics knowledge (PK) task, and a content-general transitive inference (TI) task while in the MRI scanner (**Figure 1**). The FCI is a reliable^51^ and widely used^52^ test of conceptual understanding in Newtonian Physics^50^ whose adaptation for and implementation in the MRI environment has been described in detail elsewhere^12^. FCI questions asked students to reason through a set of causal physical scenarios by forcing them to choose between a single correct answer and multiple commonsense alternatives based on documented and persistent confusions introductory physics students commonly hold. Control questions for the FCI task presented similar everyday physical scenarios but tested students on basic reading comprehension and shape identification rather than formal physics content. In contrast, the PK task is a block-design task designed to measure physics-based content knowledge. Students answered physics questions (e.g., *What is the value of the acceleration due to gravity on Earth?* with answer choices such as *9*.*81 m/s^2, 15 kg, 10 liters*, and *11 ft/s^2*), while control questions^53^ tested students’ general knowledge retrieval (e.g., *What is the tallest mountain in the world?* with answer choices such as *Mount Rushmore, Rainier Mountain, Mount Everest*, and *Mount Logan*). Finally, the TI task is a fast event-related paradigm adapted from a canonical deductive reasoning task designed to assess general reasoning ability^54,55^. Students viewed sequential relational statements (e.g., *The Fork is to the left of the Plate* and *The Fork is to the right of the Cup*) followed by a putative conclusion (e.g., *The Cup is to the left of the Plate?*) and indicated if the conclusion logically followed from the premises. Control TI questions for were of similar but illogical form (e.g., *The Fork is to the left of the Plate* and *The Fork is to the right of the Cup* followed by the non-logical conclusion *The Bowl is to the left of the Saucer?*). Schematics of timing and trial information for all tasks are provided in **Supplementary Figure 4**.

### fMRI acquisition and pre-processing

Data were collected on a General Electric 3T Healthcare Discovery 750w MRI scanner utilizing a 32-channel phased-array head radio frequency coil located at the University of Miami. Functional images were acquired obliquely using an interleaved gradient echo EPI pulse sequence (TR/TE = 2000/30ms, flip angle = 75°, field of view = 220×220mm, matrix size = 64×64, voxel dimensions = 3.4×3.4×3.4mm, 42 axial oblique slices). High-resolution T1-weighted sagittal 3D FSPGR BRAVO sequences were also collected with 186 contiguous sagittal slices (TI = 650ms, bandwidth = 25.0kHz, flip angle = 12°, FOV = 256×256mm, and slice thickness = 1.0mm) for anatomical reference. Pre-processing was performed in FSL and AFNI in which functional images were skull stripped, motion corrected, high-pass filtered (110s), and co-registered with structural volumes via a 12-degree-of-freedom affine transformation. Images were then resampled into MNI152 space and spatially smoothed (5mm FWHM Gaussian kernel). TRs with excessive motion (including one frame before and two frames after^56^) were scrubbed if they met or exceeded a threshold of 0.35mm framewise displacement^12^. Additionally, due to the relatively long trials of the FCI task, FCI runs containing excessive motion (≥33% of within-block motion) were discarded from the analysis, resulting in the omission of three runs from two individuals.

### Statistical analyses

General linear model analyses were performed in FSL using FEAT. FCI and PK task and control questions were modeled as blocks from question onset to the onset of a concluding fixation, while TI task and control questions were modeled as events occurring at the halfway point between the onset of the conclusion statement and the individual’s button press. All timing files were convolved with a hemodynamic response function and first temporal derivatives of each convolved regressor were included to account for offsets in peak BOLD response. Six motion parameters (translations and rotations) were included as nuisance regressors in all analyses. The contrasts FCI > Control, PK > Control, and TI > Control were modeled within-subject and group-level activation maps were generated and thresholded with a cluster defining threshold (CDT) of *P* < 0.001 and a cluster extent threshold (CET) of *P* < 0.05 (FWE corrected). Maps were computed by session individually, then class, sex, and longitudinal changes were assessed using a three-way fixed effects ANOVA to identify regions more engaged during task within one group relative to another, at the end relative to the beginning of the semester, and to test for significant interactions between class, sex, and time.

## DATA AVAILABILITY STATEMENT

A GitHub repository was created at https://github.com/NBCLab/PhysicsLearning/tree/master/learning to archive the source files for this study, including data analysis processing scripts and behavioral data. The network results for the FCI, PK, and TI tasks for all time points, contrasts across time, and interactions are available via NeuroVault at https://neurovault.org/collections/5939/.

## ACKNOWLEDGMENTS

Primary funding for this project was provided by NSF REAL DRL-1420627; additional support was provided by NSF 1631325, NIH R01 DA041353, NIH U01 DA041156, NSF CNS 1532061, NIH K01DA037819, NIH U54MD012393, and the FIU Graduate School Dissertation Year Fellowships. Thanks to Karina Falcone, Rosario Pintos Lobo, and Camila Uzcategui for their assistance with data collection, to Dr. Melanie Stollstorf for providing TI task stimuli, and to Dr. Jeremy Elman for providing generally known fact question that were drawn from for the PK task. Additional thanks to the FIU Instructional & Research Computing Center (IRCC, http://ircc.fiu.edu) for providing the HPC and computing resources that contributed to the research results reported within this paper, and to the Department of Psychology of the University of Miami for providing access to their MRI scanner. Lastly, special thanks to the FIU undergraduate students who volunteered, participated, and contributed to this project.

## AUTHOR CONTRIBUTIONS

ARL, EB, SMP, MTS, RWL conceived and designed the project. JEB, KLB, and ERB acquired the fMRI data, and JEB, MCR, TS, and KLB contributed scripts and pipelines. JEB performed all data analyses. JEB and ARL drafted the paper, and all authors contributed to the revisions and approved the final version.

