## Supplemental Information for "Sex and pedagogy influences in physics learning-related reorganization of brain activation"

### SUPPLEMENTAL METHODS

**Study Timeline.** Data were collected across six 15-week academic semesters from students enrolled in 21 separate University-level introductory physics course sections. Pre-instruction imaging sessions were acquired during the period starting one week prior to the first class meeting and ending no more than four weeks into the academic semester, before the first physics course exam. Post-instruction imaging sessions were conducted after the final exam of each physics course. In most cases this meant that students underwent MRI scanning in the two weeks following the University's final exam week. However, in some cases (e.g., due to conflicts in individual's post-semester travel schedules) students were unable to complete post-instruction MRI scanning after the semester ended. Thus, a total of 15 individuals completed post-instruction MRI scanning during finals week. For these sessions the study team attempted to schedule the MRI scan after the completion of the student's full set of finals exams, resulting in 13 students who completing their post-instruction MRI scan on the last day of finals week, after all exams had concluded.

**Gender and Sex.** In everyday language and in scientific research, the terms "gender" and "sex" are often used interchangeably. The two concepts, however, are not the same. Sex is biologically determined and refers to whether an individual possesses two X chromosomes or an XY pair. The terms "Male" and "Female" describe biological sex, and widespread confusion about their specific biological meaning results in their frequent misuse. Gender, on the other hand, describes a social construct that refers to either societal expectations or behaviors traditionally ascribed to women and men (i.e., gender roles), or how someone self-identifies as a woman, man, or non-binary (i.e., gender identity). Many individuals who are biologically male or female also identify with being a man or a woman; however, this is not always the case and assuming so is both inappropriate and misrepresentative. In this study, we did not explore the neurobiological influences associated with gender in physics classrooms, although this question is most certainly important and extremely relevant to the current work. During data collection, we relied on demographics collected by Florida International University at the time of each student's initial enrollment in their classes. In this demographics assessment students indicated whether or not they were male or female. Thus, our study reports on sex influences in physics-related brain function. Additional measures on gender identity were, regrettably, not collected. Student gender is most certainly relevant, and indeed highly influential, to the topics and findings discussed in this study. Future work will place additional focus on the influences gender identity, societal expectations, and related gendered factors may have on brain function.

**Course Grades.** All students were graded on the Florida International University 4-point system in which course grades are determined on the following numerical scale: an A is 4.00, an A- is 3.67, a B+ is 3.33, a B is 3.00, a B- is 2.67, a C+ is 2.33, a C is 2.00, a C- is 1.67, a D+ is 1.33, a D is 1.00, a D- is 0.67, and an F is 0.00 points per credit hour. The distribution of course grades for the students who took part in this study are provided in **Supplementary Figure 1**.

**Meta-analytic Functional Decoding Using Neurosynth.** Probabilistically leveraging the wider neuroimaging literature to guide the interpretation of experimental results is an arguably more objective approach than speculating about the meaning of results based on only a limited set of citations brought into the discussion. Unsurprisingly, many researchers have begun to adopt this technique and increasingly sophisticated probabilistic tools and frameworks for 'decoding' brain activity have been developed[1-3]. Neurosynth (http://neurosynth.org)[3] is a large database (>11,000 neuroimaging studies) of mappings between neural states and cognitive processes that can be used to perform quantitative reverse inference. Neurosynth uses automated text mining algorithms to access published neuroimaging studies and extract meaningful words (or 'terms') from the paper's abstract along with peak activation coordinate results as reported in the paper's tables. Terms are thought to represent cognitive and/or psychological processes that are linked to the activation coordinates, although some terms may be ambiguous to interpret due to de-contextualization or multiple terms represent similar mental states. In this paper, we make use of a Generalized Correspondence Latent Dirichlet Allocation (GC-LDA) machine learning algorithm[1,2] which was trained on the Neurosynth database to determine likely sets of underlying mental functions that are associated with our observed FCI-related brain activity. GC-LDA is a data reduction technique that was used to detect a set of latent and semantically coherent 'topics' that are present in the Neurosynth database. Each topic consists of probability distributions of spatial activation coordinates and semantically linked terms. Thus, topics form highly interpretable collections of functional brain regions linked with mental processes that can then be used to decode whole-brain activation patterns[1]. In this study we fed in the unthresholded z-statistic brain activation maps into the GC-LDA topic model using 200 distinct topics to determine a rank ordered lists of terms that, according the latent topics present in the Neurosynth database, are most associated with the input pattern of brain activity observed during the FCI. We did this to form a data-driven text-based representation of each observed activation pattern so as to objectively guide our interpretation of results.

### SUPPLEMENTAL FIGURES

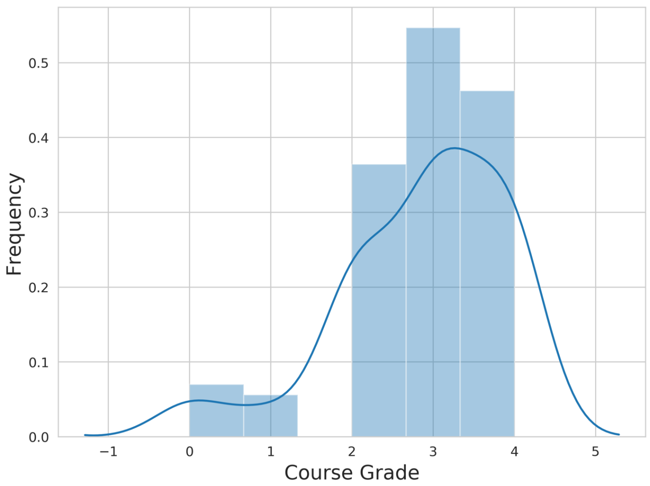


**Supplementary Figure 1.** Student Course Grades. *Distribution of course grades received by students who took part in the study.*

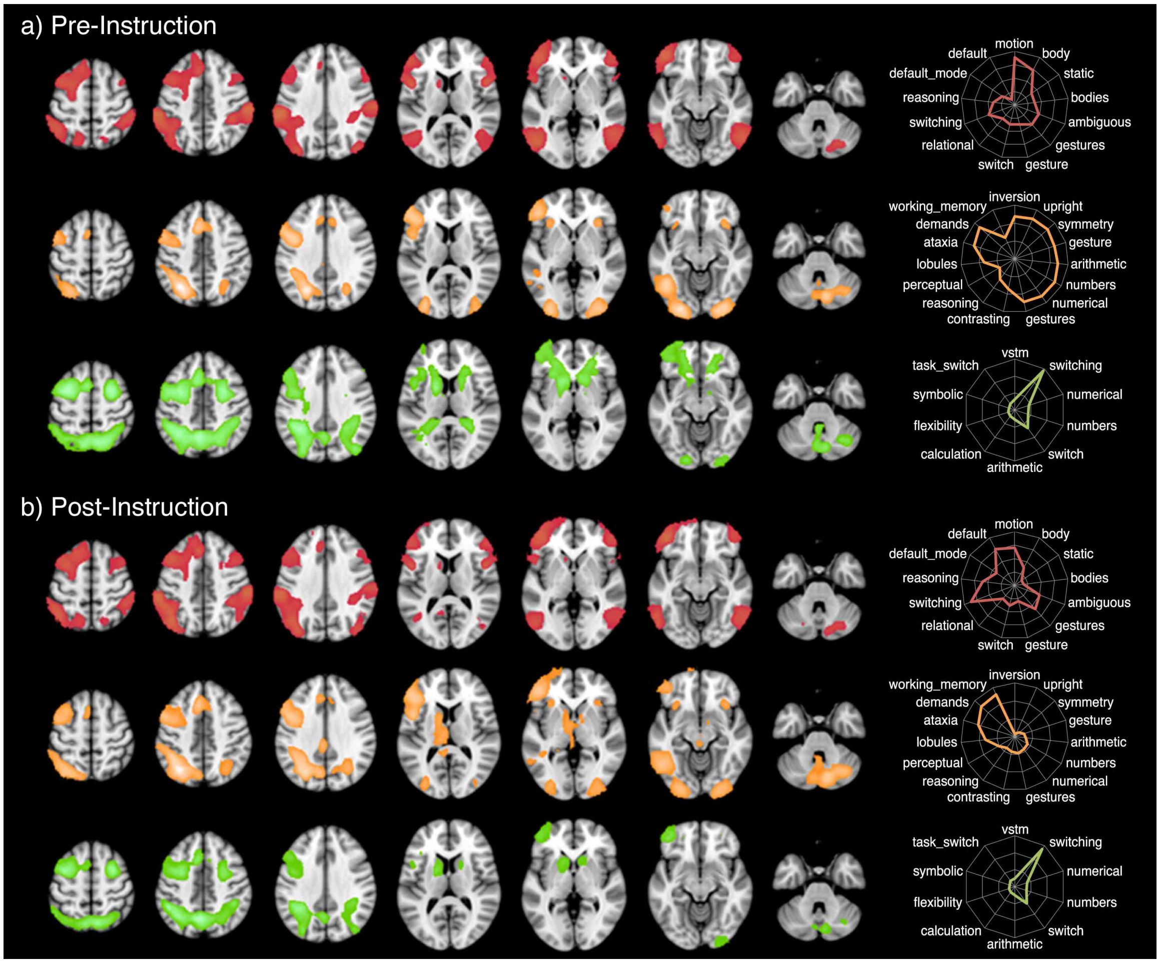


**Supplementary Figure 2.** Task Activations Before and After Instruction. *a) Pre- and b) post-instruction activation maps for all students and all tasks, including FCI > Control (rose), PK > Control (orange), and TI > Control (green). Radar plots (arbitrary units) give GC-LDA functional decoding*[2] *of each continuous z-map. Activation maps were thresholded using a cluster defining threshold of $P < 0.001$ and a cluster extent threshold of $P < 0.05$, FWE corrected. Radar plots display the union of the top 10 Neurosynth decoding terms within each task for the pre- and post-instruction z-maps (see Table S2 for Neurosynth terms and weights for each contrast). If any term produced by the decoder was unmatched (e.g., was ranked ≥10 within the paradigm for one z-map but <10 for the other) then the lesser-ranked term was included in the paradigm's radar plot for visualization.*

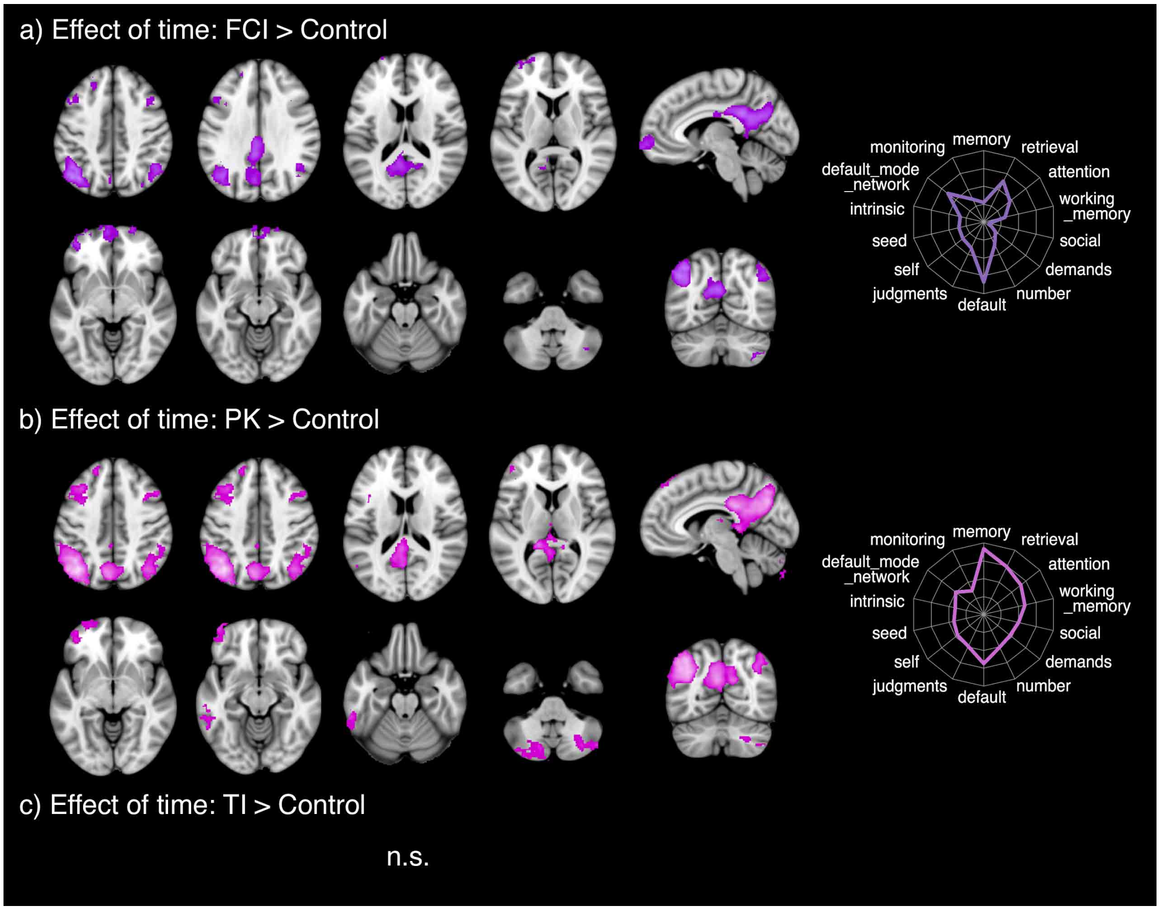


**Supplemental Figure 3.** Changes in Task Activation Across Instruction. *a) Post > Pre: FCI > Control (purple), b) Post > Pre: PK > Control (pink), and c) Post > Pre: TI > Control (no significant activity was observed). Activation maps were thresholded using a cluster defining threshold of P < 0.001 and a cluster extent threshold of P < 0.05, FWE corrected. Color-coded radar plots (arbitrary units) show GC-LDA[2] meta-analytic functional decoding for each Post > Pre condition map (see Table S4 for functional terms and weights for each contrast).*

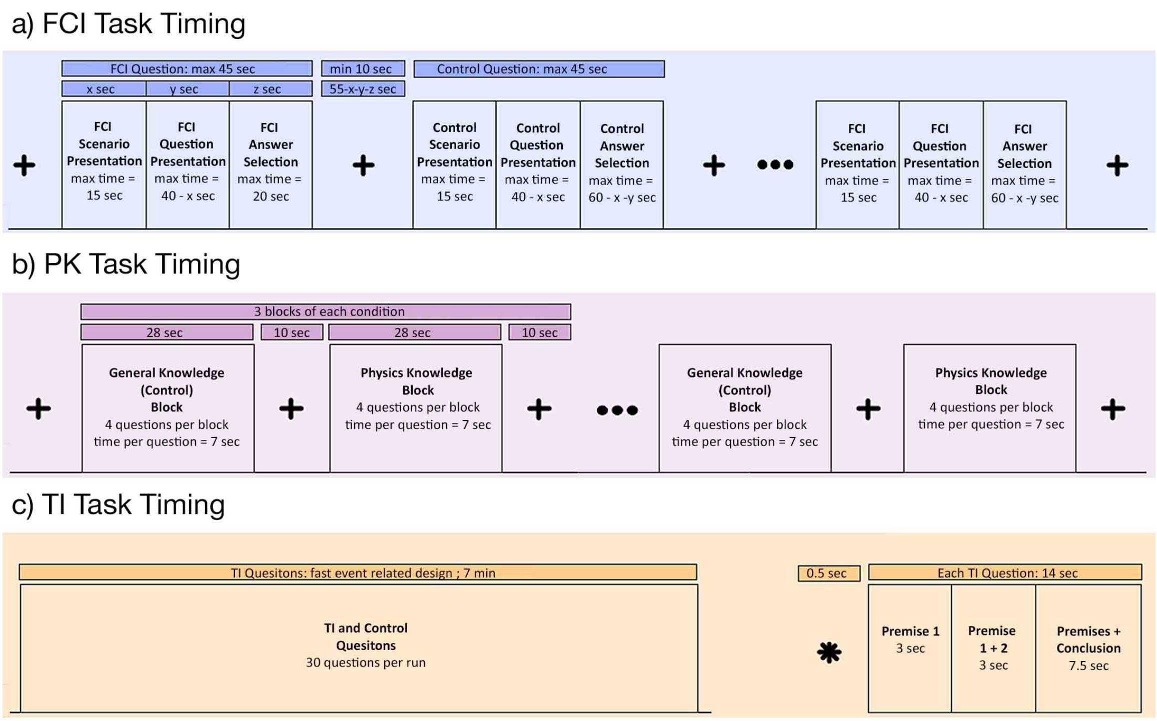


**Supplemental Figure 4.** Schematics of Timing and Trial Information for the a) FCI, b) PK, and c) TI Task Paradigms.

### SUPPLEMENTAL TABLES

**Supplemental Table 1.** Brain Activation Results at Pre- and Post-Instruction. *Center of mass coordinates associated with brain activation maps for FCI > Control, PK > Control, and TI > Control at (a-c) pre- and (d-f) post-instruction stages. Cluster region labels are based off those reported by the IBASPM116 Human Brain Atlas. Coordinates are reported in MNI space.*

| **a) Pre-Instruction: FCI > Control** | | | | | | | |
|---|---|---|---|---|---|---|---|
| Cluster | Hemisphere | Center of Mass (MNI space) | | | Cluster Extent ($mm^3$) | Mean Z Score | Labels |
| | | X | Y | Z | | | |
| 1 | B | -36 | 26 | 26 | 79848 | 5.892 | White Matter, Frontal_Mid_L, Frontal_Inf_Tri_L, Frontal_Sup_L, Frontal_Sup_Medial_L, Frontal_Inf_Oper_L, Precentral_L, Frontal_Inf_Orb_L, Supp_Motor_Area_L, Frontal_Mid_Orb_L, Rolandic_Oper_L, Frontal_Sup_Orb_L, Frontal_Sup_Medial_R, Cingulum_Mid_L, Temporal_Pole_Sup_L, Supp_Motor_Area_R |
| 2 | L | -48 | -54 | 28 | 72016 | 5.972 | White Matter, Parietal_Inf_L, Temporal_Mid_L, Occipital_Mid_L, SupraMarginal_L, Angular_L, Temporal_Inf_L, Parietal_Sup_L, Occipital_Inf_L, Precuneus_L, Postcentral_L, Temporal_Sup_L, Occipital_Sup_L |
| 3 | R | 50 | -62 | 0 | 34416 | 5.706 | White Matter, Temporal_Mid_R, Temporal_Inf_R, Occipital_Mid_R, Fusiform_R, Occipital_Inf_R, Angular_R, Cerebelum_Crus1_R, Cerebelum_6_R |
| 4 | R | 50 | 28 | 12 | 27968 | 4.904 | White Matter, Frontal_Inf_Tri_R, Frontal_Inf_Oper_R, Frontal_Mid_R, Frontal_Inf_Orb_R, Frontal_Mid_Orb_R, Precentral_R, Rolandic_Oper_R, Insula_R |
| 5 | R | 50 | -36 | 44 | 26712 | 5.935 | White Matter, SupraMarginal_R, Parietal_Inf_R, Postcentral_R, Parietal_Sup_R, Precuneus_R, Angular_R |
| 6 | R | 32 | -72 | -44 | 11160 | 5.354 | |
| 7 | L | -12 | 10 | 8 | 1304 | 4.445 | Caudate_L, White Matter |
| **b) Pre-Instruction: PK > Control** | | | | | | | |
| 1 | L | -34 | -68 | 14 | 66952 | 6.249 | White Matter, Occipital_Mid_L, Parietal_Inf_L, Temporal_Inf_L, |

| | | | | | | | |
|---|---|---|---|---|---|---|---|
| | | | | | | | Parietal_Sup_L, Occipital_Inf_L, Fusiform_L, Temporal_Mid_L, Lingual_L, Angular_L, Occipital_Sup_L, Precuneus_L, Cerebelum_Crus1_L, Cuneus_L, Calcarine_L, SupraMarginal_L, Cerebelum_6_L |
| 2 | L | -46 | 18 | 22 | 50856 | 6.213 | White Matter, Frontal_Inf_Tri_L, Precentral_L, Frontal_Inf_Oper_L, Frontal_Mid_L, Frontal_Inf_Orb_L, Insula_L, Frontal_Mid_Orb_L, Rolandic_Oper_L, Postcentral_L, Frontal_Sup_L |
| 3 | B | 18 | -70 | -42 | 26544 | 5.051 | Cerebro-Spinal Fluid |
| 4 | R | 30 | -84 | 4 | 22808 | 5.759 | White Matter, Occipital_Mid_R, Occipital_Inf_R, Lingual_R, Occipital_Sup_R, Calcarine_R, Angular_R, Fusiform_R, Parietal_Sup_R, Parietal_Inf_R, Cerebelum_Crus1_R, Precuneus_R, Cuneus_R |
| 5 | B | 0 | 22 | 40 | 10760 | 5.227 | Supp_Motor_Area_L, Frontal_Sup_Medial_L, Cingulum_Mid_R, Cingulum_Ant_R, Cingulum_Mid_L, Supp_Motor_Area_R, Frontal_Sup_L, Cingulum_Ant_L, Frontal_Sup_Medial_R, White Matter |
| 6 | B | -4 | -24 | 24 | 5488 | 4.667 | White Matter, Cerebro-Spinal Fluid, Cingulum_Post_L, Cingulum_Mid_L, Cingulum_Mid_R, Cingulum_Ant_L |
| 7 | R | 34 | 24 | -6 | 2864 | 5.059 | White Matter, Insula_R, Frontal_Inf_Orb_R, Frontal_Inf_Tri_R, Putamen_R |
| **c) Pre-Instruction: TI > Control** | | | | | | | |
| 1 | B | -12 | -22 | 34 | 186568 | 5.279 | White Matter, Precentral_L, Frontal_Mid_L, Parietal_Inf_L, Precuneus_L, Precuneus_R, Frontal_Inf_Tri_L, Parietal_Sup_L, Caudate_L, Parietal_Sup_R, Occipital_Mid_L, Frontal_Mid_Orb_L, Frontal_Inf_Oper_L, Frontal_Sup_L, Supp_Motor_Area_L, Parietal_Inf_R, Frontal_Mid_R, Frontal_Sup_R, Occipital_Mid_R, Angular_L, Putamen_L, Angular_R, Frontal_Sup_Orb_L, Occipital_Sup_R, Frontal_Inf_Orb_L, Precentral_R, Frontal_Sup_Medial_L, Insula_L, Occipital_Sup_L, Pallidum_L, SupraMarginal_R, Postcentral_L, Temporal_Mid_L, Supp_Motor_Area_R, SupraMarginal_L, Cingulum_Mid_L, Cingulum_Mid_R, Rolandic_Oper_L, Rectus_L, Cuneus_R, Cingulum_Ant_L, |

| | | | | | | | |
|---|---|---|---|---|---|---|---|
| | | | | | | | Hippocampus_L, Calcarine_R, Olfactory_L, Cingulum_Post_L, Frontal_Mid_Orb_L, Cingulum_Post_R, Hippocampus_R, Postcentral_R, Cerebro-Spinal Fluid |
| 2 | R | 20 | 26 | 2 | 19008 | 4.213 | White Matter, Caudate_R, Frontal_Sup_Orb_R, Putamen_R, Frontal_Mid_Orb_R, Pallidum_R, Rectus_R, Frontal_Inf_Tri_R, Frontal_Inf_Orb_R, Olfactory_R, Frontal_Mid_Orb_R, Insula_R, Frontal_Mid_R, Cingulum_Ant_R, Cerebro-Spinal Fluid |
| 3 | B | 2 | -62 | -32 | 7104 | 4.339 | Cerebro-Spinal Fluid |
| 4 | R | 34 | -64 | -38 | 3120 | 4.667 | |
| 5 | R | 28 | -92 | -12 | 1904 | 3.452 | White Matter, Occipital_Inf_R, Lingual_R, Fusiform_R |
| 6 | L | -18 | -92 | -10 | 1272 | 3.904 | White Matter, Lingual_L, Occipital_Inf_L, Occipital_Mid_L, Calcarine_L, Fusiform_L |
| 7 | R | 32 | 52 | -2 | 16 | 3.134 | White Matter |
| **d) Post-Instruction: FCI > Control** | | | | | | | |
| 1 | B | -34 | 30 | 26 | 85072 | 5.676 | White Matter, Frontal_Mid_L, Frontal_Sup_L, Frontal_Inf_Tri_L, Frontal_Sup_Medial_L, Precentral_L, Frontal_Inf_Oper_L, Frontal_Inf_Orb_L, Frontal_Mid_Orb_L, Supp_Motor_Area_L, Frontal_Sup_Orb_L, Frontal_Mid_Orb_L, Rolandic_Oper_L, Cingulum_Mid_L, Cingulum_Ant_L, Frontal_Sup_Medial_R, Supp_Motor_Area_R, Temporal_Pole_Sup_L |
| 2 | R | 50 | -50 | 26 | 57088 | 5.222 | White Matter, Temporal_Inf_R, SupraMarginal_R, Parietal_Inf_R, Temporal_Mid_R, Occipital_Mid_R, Parietal_Sup_R, Angular_R, Postcentral_R, Occipital_Inf_R, Fusiform_R, Precuneus_R, Cerebelum_Crus1_R, Occipital_Sup_R, Cerebelum_6_R |
| 3 | L | -46 | -54 | 42 | 49632 | 6.386 | White Matter, Parietal_Inf_L, Angular_L, SupraMarginal_L, Parietal_Sup_L, Occipital_Mid_L, Precuneus_L, Postcentral_L, Occipital_Sup_L, Temporal_Sup_L, Temporal_Mid_L |
| 4 | R | 46 | 28 | 18 | 36576 | 4.746 | White Matter, Frontal_Mid_R, Frontal_Inf_Tri_R, Frontal_Inf_Oper_R, Frontal_Mid_Orb_R, Frontal_Inf_Orb_R, Frontal_Sup_R, Precentral_R, Rolandic_Oper_R, Insula_R |

| | | | | | | | |
|---|---|---|---|---|---|---|---|
| 5 | L | -54 | -58 | -8 | 17864 | 5.243 | White Matter, Temporal_Inf_L, Temporal_Mid_L, Occipital_Inf_L, Occipital_Mid_L, Cerebelum_Crus1_L |
| 6 | R | 28 | -70 | -44 | 13744 | 5.257 | |
| 7 | L | -32 | -74 | -52 | 6120 | 4.148 | |
| 8 | L | -8 | -56 | 16 | 1680 | 3.666 | White Matter, Precuneus_L, Calcarine_L, Cingulum_Post_L, Cuneus_L |
| 9 | L | -12 | 10 | 8 | 1392 | 3.997 | Caudate_L, White Matter |
| **e) Post-Instruction: PK > Control** | | | | | | | |
| 1 | B | -2 | -54 | 28 | 38936 | 4.242 | White Matter, Precuneus_L, Precuneus_R, Cingulum_Post_L, Cuneus_L, Cingulum_Mid_L, Cingulum_Mid_R, Cingulum_Post_R, Calcarine_L, Cuneus_R, Hippocampus_L, Thalamus_L, Vermis_4_5, Occipital_Sup_L, Parietal_Sup_L, Cerebelum_4_5_L, Lingual_L, Calcarine_R, Parietal_Sup_R, Lingual_R, Thalamus_R, Cerebro-Spinal Fluid |
| 2 | L | -42 | -62 | 38 | 30520 | 4.683 | White Matter, Angular_L, Parietal_Inf_L, Occipital_Mid_L, Parietal_Sup_L, Temporal_Mid_L, SupraMarginal_L, Occipital_Sup_L |
| 3 | B | 0 | -74 | -44 | 16056 | 3.502 | |
| 4 | B | -32 | 20 | 46 | 15520 | 3.766 | White Matter, Frontal_Mid_L, Frontal_Sup_Medial_L, Frontal_Sup_L, Precentral_L, Frontal_Inf_Oper_L, Frontal_Inf_Tri_L, Frontal_Sup_Medial_R |
| 5 | R | 44 | -60 | 44 | 10888 | 3.876 | White Matter, Angular_R, Parietal_Inf_R, Occipital_Mid_R, SupraMarginal_R, Parietal_Sup_R, Occipital_Sup_R |
| 6 | L | -38 | 52 | -4 | 6384 | 3.595 | White Matter, Frontal_Mid_Orb_L, Frontal_Inf_Tri_L, Frontal_Mid_L, Frontal_Sup_Orb_L, Frontal_Inf_Orb_L, Frontal_Sup_L |
| 7 | R | 40 | 14 | 48 | 5104 | 3.890 | White Matter, Frontal_Mid_R, Frontal_Inf_Oper_R, Precentral_R, Frontal_Inf_Tri_R |
| 8 | L | -58 | -40 | -16 | 3672 | 3.491 | White Matter, Temporal_Inf_L, Temporal_Mid_L |
| 9 | B | -2 | -92 | -24 | 448 | 3.262 | Cerebelum_Crus2_L |
| 10 | R | 12 | -42 | 10 | 200 | 3.528 | White Matter |
| 11 | R | 16 | -36 | 8 | 16 | 3.221 | White Matter |
| 12 | R | 50 | 28 | 36 | 16 | 3.160 | Frontal_Mid_R, White Matter |
| 13 | R | 52 | 26 | 34 | 8 | 3.218 | White Matter |
| **f) Post-Instruction: TI > Control** | | | | | | | |
| 1 | B | -8 | -58 | 42 | 72656 | 5.541 | White Matter, Parietal_Inf_L, Precuneus_L, Precuneus_R, |

| | | | | | | | |
|---|---|---|---|---|---|---|---|
| | | | | | | | Parietal_Sup_L, Occipital_Mid_L, Parietal_Sup_R, Parietal_Inf_R, Angular_L, Occipital_Mid_R, Angular_R, Occipital_Sup_R, SupraMarginal_R, Occipital_Sup_L, SupraMarginal_L, Temporal_Mid_L, Cuneus_R, Cingulum_Post_L, Cingulum_Post_R, Cerebro-Spinal Fluid |
| 2 | B | -34 | 10 | 42 | 48176 | 5.051 | White Matter, Precentral_L, Frontal_Mid_L, Frontal_Inf_Tri_L, Frontal_Inf_Oper_L, Frontal_Sup_L, Supp_Motor_Area_L, Frontal_Sup_Medial_L, Supp_Motor_Area_R, Rolandic_Oper_L, Cingulum_Mid_L, Postcentral_L, Insula_L, Cingulum_Mid_R |
| 3 | L | -38 | 52 | -8 | 13760 | 4.650 | White Matter, Frontal_Mid_Orb_L, Frontal_Mid_L, Frontal_Inf_Orb_L, Frontal_Sup_Orb_L, Frontal_Inf_Tri_L, Frontal_Sup_L |
| 4 | R | 28 | 2 | 50 | 11576 | 5.600 | White Matter, Frontal_Mid_R, Frontal_Sup_R, Precentral_R |
| 5 | L | -14 | 8 | 8 | 8856 | 4.418 | Caudate_L, Putamen_L, Pallidum_L, White Matter, Cerebro-Spinal Fluid |
| 6 | R | 14 | 10 | 8 | 4040 | 3.822 | Caudate_R, Putamen_R, Pallidum_R, Olfactory_R, White Matter, Cerebro-Spinal Fluid |
| 7 | R | 28 | -92 | -12 | 2896 | 3.869 | White Matter, Occipital_Inf_R, Lingual_R, Fusiform_R, Calcarine_R |
| 8 | R | 32 | -64 | -38 | 2616 | 4.509 | |
| 9 | B | 6 | -76 | -34 | 2552 | 4.247 | |
| 10 | R | 24 | 50 | -20 | 1424 | 3.652 | Frontal_Mid_Orb_R, Frontal_Sup_Orb_R, White Matter |

**Supplemental Table 2.** Meta-analytic Decoding Results at Pre- and Post-Instruction. *Meta-analytic functional decoding was performed on unthresholded z-statistic maps for (a) Pre-instruction: FCI > Control, (b) Pre-instruction: PK > Control, (c) Pre-instruction: TI > Control, (d) Post-instruction: FCI > Control, (e) Post-instruction: PK > Control, and (e) Post-instruction: TI > Control. Decoding was performed with a 200-topic GC-LDA[2] topic model trained on the Neurosynth database. The top 10 terms returned for each map are provided alongside their associated correlation values. Note these weights depend on the input volume and therefore are meaningful as relative values within each map, however weights should not be assigned an absolute interpretation across images[2].*

| **a) Pre-instruction: FCI > Control** | |
|---|---|
| **Term** | **Weight** |
| motion | 633.7252146 |
| body | 478.3801832 |
| switching | 267.1527428 |
| gestures | 233.9310515 |
| ambiguous | 225.5804697 |
| reasoning | 153.3920201 |
| static | 126.1253032 |
| bodies | 123.1749244 |
| switch | 118.3710158 |
| gesture | 117.8680682 |
| **b) Pre-instruction: PK > Control** | |
| **Term** | **Weight** |
| demands | 81.68425191 |
| numbers | 74.32884743 |
| numerical | 73.4534514 |
| symmetry | 60.42453932 |
| gestures | 58.331436 |
| upright | 58.30865644 |
| arithmetic | 54.99401087 |
| ataxia | 49.61921856 |
| inversion | 47.62903342 |
| gesture | 43.12472325 |
| **c) Pre-instruction: TI > Control** | |
| **Term** | **Weight** |
| switching | 573.428806 |
| switch | 262.5824401 |
| numerical | 174.3624206 |
| numbers | 164.7832427 |
| vstm | 126.2531562 |
| arithmetic | 112.7522074 |
| calculation | 90.72827503 |
| task_switching | 89.26145701 |
| flexibility | 80.30573393 |
| symbolic | 69.79990286 |
| **d) Post-instruction: FCI > Control** | |
| **Term** | **Weight** |
| switching | 309.71084 |
| default | 276.1635173 |
| motion | 252.5753468 |
| reasoning | 214.5672386 |
| gestures | 211.2976516 |

| | |
|---|---|
| ambiguous | 180.1752178 |
| default_mode | 147.8986879 |
| switch | 137.3589833 |
| body | 135.9365735 |
| relational | 117.1724557 |
| **e) Post-instruction: PK > Control** | |
| **Term** | **Weight** |
| working_memory | 463.7304765 |
| demands | 453.0719902 |
| ataxia | 386.9300548 |
| lobules | 297.2108844 |
| perceptual | 187.2265326 |
| gestures | 169.3008325 |
| contrasting | 154.8410312 |
| numerical | 153.667509 |
| numbers | 147.7564221 |
| reasoning | 138.9909681 |
| **f) Post-instruction: TI > Control** | |
| **Term** | **Weight** |
| switching | 547.9865903 |
| switch | 243.6652172 |
| numerical | 151.5781711 |
| numbers | 145.1158657 |
| arithmetic | 101.4927589 |
| vstm | 98.64800143 |
| task_switching | 85.15796113 |
| calculation | 79.59715502 |
| flexibility | 72.23492967 |
| symbolic | 62.86652072 |

**Supplemental Table 3.** Brain Activation Results for Post- > Pre-Instruction. *Center of mass coordinates associated with brain activation maps for a) Post > Pre-Instruction: FCI > Control and b) Post > Pre-Instruction: PK > Control. Post > Pre-Instruction: TI > Control was tested but no significant effect of time was detected. Cluster region labels are based off those reported by the IBASPM116 Human Brain Atlas. Coordinates are reported in MNI space.*

| Cluster | Hemisphere | Center of Mass (MNI space) | | | Cluster Extent ($mm^3$) | Mean Z Score | Labels |
|---|---|---|---|---|---|---|---|
| | | X | Y | Z | | | |
| **a) Post > Pre Instruction: FCI > Control** | | | | | | | |
| 1 | B | -4 | -52 | 24 | 19608 | 4.038 | White Matter, Precuneus_L, Cingulum_Post_L, Cuneus_L, Precuneus_R, Calcarine_L, Cingulum_Mid_L, Cingulum_Mid_R, Cingulum_Post_R, Calcarine_R, Occipital_Sup_L, Cuneus_R, Lingual_L, Cerebelum_4_5_L |
| 2 | L | -42 | -64 | 42 | 13096 | 4.339 | White Matter, Angular_L, Parietal_Inf_L, Occipital_Mid_L, Parietal_Sup_L, Occipital_Sup_L, SupraMarginal_L |
| 3 | B | -14 | 62 | -2 | 9552 | 3.587 | Frontal_Mid_Orb_L, Frontal_Mid_L, Frontal_Sup_L, Frontal_Mid_Orb_L, Frontal_Mid_Orb_R, Frontal_Sup_Medial_L, Frontal_Sup_Orb_L, Frontal_Sup_Orb_R, Frontal_Sup_R, Frontal_Mid_Orb_R, Frontal_Inf_Tri_L, White Matter |
| 4 | R | 46 | -60 | 42 | 5144 | 3.772 | White Matter, Angular_R, Parietal_Inf_R, Occipital_Sup_R, Parietal_Sup_R, Occipital_Mid_R |
| 5 | L | -44 | 18 | 38 | 2168 | 3.539 | White Matter, Frontal_Mid_L, Frontal_Inf_Oper_L, Frontal_Inf_Tri_L |
| 6 | R | 40 | 16 | 46 | 1856 | 3.469 | White Matter, Frontal_Mid_R, Frontal_Inf_Oper_R |
| 7 | R | 40 | -62 | -48 | 1200 | 3.693 | |
| 8 | L | -20 | 40 | 40 | 1168 | 3.392 | White Matter, Frontal_Sup_L, Frontal_Mid_L |
| **b) Post > Pre Instruction: PK > Control** | | | | | | | |
| 1 | B | -2 | -52 | 28 | 37120 | 4.215 | White Matter, Precuneus_L, Precuneus_R, Cingulum_Post_L, Cuneus_L, Cingulum_Mid_L, Cingulum_Post_R, Cingulum_Mid_R, Calcarine_L, Thalamus_L, Hippocampus_L, Cuneus_R, Vermis_4_5, Occipital_Sup_L, Cerebelum_4_5_L, Lingual_L, Parietal_Sup_L, Calcarine_R, Parietal_Sup_R, Lingual_R, Thalamus_R, |

| | | | | | | | |
|---|---|---|---|---|---|---|---|
| | | | | | | | Cerebro-Spinal Fluid |
| 2 | L | -42 | -62 | 38 | 29584 | 4.688 | White Matter, Angular_L, Parietal_Inf_L, Occipital_Mid_L, Parietal_Sup_L, Temporal_Mid_L, SupraMarginal_L, Occipital_Sup_L |
| 3 | B | -32 | 18 | 46 | 13864 | 3.794 | White Matter, Frontal_Mid_L, Frontal_Sup_L, Precentral_L, Frontal_Sup_Medial_L, Frontal_Inf_Oper_L, Frontal_Inf_Tri_L, Frontal_Sup_Medial_R |
| 4 | R | 44 | -60 | 44 | 10232 | 3.829 | White Matter, Angular_R, Parietal_Inf_R, Occipital_Mid_R, SupraMarginal_R, Parietal_Sup_R, Occipital_Sup_R |
| 5 | L | -26 | -78 | -44 | 8072 | 3.461 | |
| 6 | L | -38 | 52 | -4 | 5904 | 3.583 | White Matter, Frontal_Mid_Orb_L, Frontal_Inf_Tri_L, Frontal_Mid_L, Frontal_Sup_Orb_L, Frontal_Inf_Orb_L, Frontal_Sup_L |
| 7 | R | 32 | -70 | -42 | 4936 | 3.491 | |
| 8 | R | 40 | 14 | 48 | 4736 | 3.872 | White Matter, Frontal_Mid_R, Frontal_Inf_Oper_R, Precentral_R, Frontal_Inf_Tri_R |
| 9 | L | -58 | -40 | -16 | 3656 | 3.557 | White Matter, Temporal_Inf_L, Temporal_Mid_L |
| 10 | R | 12 | -42 | 10 | 200 | 3.517 | White Matter |
| 11 | R | 8 | -74 | 58 | 48 | 3.243 | |
| 12 | R | 16 | -36 | 8 | 16 | 3.239 | White Matter |
| 13 | L | -40 | 34 | 8 | 8 | 3.100 | White Matter |

**Supplemental Table 4.** Meta-analytic Decoding Results for Post- > Pre-Instruction. *Meta-analytic functional decoding was performed on unthresholded z-statistic maps for (a) Pre > Post-instruction: FCI > Control and (b) Pre > Post-instruction: PK > Control. Decoding was performed with a 200-topic GC-LDA[2] topic model trained on the Neurosynth database. The top 10 terms returned for each map are provided alongside their associated correlation values. Note these weights depend on the input volume and therefore are meaningful as relative values within each map, however weights should not be assigned an absolute interpretation across images[2].*

| **a) Post > Pre: FCI > Control** | |
|---|---|
| **Term** | **Weight** |
| default | 402.7355617 |
| default_mode_network | 306.1469811 |
| retrieval | 303.8934813 |
| attention | 225.7525699 |
| judgments | 191.2534107 |
| self | 179.4875854 |
| number | 172.1408199 |
| seed | 171.485595 |
| monitoring | 169.6543649 |
| intrinsic | 161.5914066 |
| **b) Post > Pre: PK > Control** | |
| **Term** | **Weight** |
| memory | 859.1002349 |
| retrieval | 645.9554217 |
| default | 583.7001215 |
| attention | 555.874132 |
| working_memory | 470.6065034 |
| number | 372.649867 |
| social | 362.5165474 |
| default_mode_network | 356.9872656 |
| judgments | 353.7635182 |
| demands | 325.5406406 |

**Supplemental Table 5.** Brain Activation Results for Sex Differences at Pre- and Post-Instruction. *Center of mass coordinates associated with brain activation maps for a) Pre-Instruction FCI > Control: Male > Female, b) Pre-Instruction FCI > Control: Female > Male, c) Post-Instruction FCI > Control: Male > Female, d) Post-Instruction FCI > Control: Female > Male, e) Pre-Instruction PK > Control: Male > Female, f) Pre-Instruction PK > Control: Female > Male, g) Post-Instruction PK > Control: Male > Female, and h) Post-Instruction PK > Control: Female > Male. Cluster region labels are based off those reported by the IBASPM116 Human Brain Atlas. Coordinates are reported in MNI space.*

| Cluster | Hemisphere | Center of Mass (MNI space) | | | Cluster Extent ($mm^3$) | Mean Z Score | Labels |
|---|---|---|---|---|---|---|---|
| **a) Pre-Instruction FCI > Control: Male > Female** | | | | | | | |
| | | X | Y | Z | | | |
| 1 | L | -38 | 32 | 18 | 18808 | 3.705 | White Matter, Frontal_Mid_L, Frontal_Inf_Orb_L, Frontal_Sup_L, Frontal_Inf_Tri_L, Frontal_Mid_Orb_L, Frontal_Sup_Medial_L, Precentral_L, Supp_Motor_Area_L |
| 2 | L | -54 | -42 | 42 | 11432 | 3.970 | Parietal_Inf_L, SupraMarginal_L, Parietal_Sup_L, Angular_L, Postcentral_L, Temporal_Sup_L, White Matter |
| 3 | R | 56 | -32 | 44 | 6344 | 3.838 | SupraMarginal_R, Parietal_Inf_R, Postcentral_R, Parietal_Sup_R, White Matter |
| 4 | R | 48 | 44 | -4 | 4792 | 3.636 | White Matter, Frontal_Inf_Tri_R, Frontal_Inf_Orb_R, Frontal_Mid_R, Frontal_Mid_Orb_R |
| 5 | R | 48 | -74 | 16 | 2064 | 3.690 | White Matter, Temporal_Mid_R, Occipital_Mid_R |
| **b) Pre-Instruction FCI > Control: Female > Male** | | | | | | | |
| 1 | B | 8 | -64 | 30 | 6480 | 3.646 | White Matter, Precuneus_R, Cuneus_R, Cuneus_L, Cingulum_Post_R, Cingulum_Mid_R, Precuneus_L |
| 2 | R | 30 | 18 | 8 | 5568 | 3.549 | White Matter, Insula_R, Caudate_R, Putamen_R, Frontal_Inf_Tri_R, Frontal_Inf_Orb_R, Frontal_Inf_Oper_R |
| 3 | R | 28 | 50 | 22 | 4304 | 3.676 | White Matter, Frontal_Mid_R, Frontal_Sup_R |
| 4 | R | 48 | -2 | 42 | 2664 | 3.627 | Precentral_R, Frontal_Mid_R, White Matter |
| 5 | B | 0 | -84 | 4 | 2512 | 3.645 | Calcarine_L, Lingual_L, Calcarine_R, Lingual_R, Cuneus_R, White Matter |
| 6 | B | 4 | 6 | 60 | 1568 | 3.557 | Supp_Motor_Area_R, Supp_Motor_Area_L, White Matter |
| **c) Post-Instruction FCI > Control: Male > Female** | | | | | | | |
| 1 | L | -54 | -46 | 40 | 8352 | 3.820 | Parietal_Inf_L, SupraMarginal_L, Angular_L, White Matter |
| 2 | L | -46 | 48 | -10 | 4120 | 3.566 | White Matter, Frontal_Mid_Orb_L, |

| | | | | | | | |
|---|---|---|---|---|---|---|---|
| | | | | | | | Frontal_Inf_Orb_L, Frontal_Mid_L, Frontal_Inf_Tri_L |
| 3 | R | 58 | -34 | 40 | 3888 | 3.683 | SupraMarginal_R, Parietal_Inf_R, Parietal_Sup_R, Postcentral_R, White Matter |
| 4 | L | -28 | 2 | 56 | 1808 | 3.780 | White Matter, Frontal_Mid_L, Precentral_L, Frontal_Sup_L |
| 5 | R | 44 | 44 | -18 | 1224 | 3.441 | White Matter, Frontal_Inf_Orb_R, Frontal_Mid_Orb_R |
| 6 | R | 46 | 50 | -4 | 336 | 3.239 | White Matter, Frontal_Mid_Orb_R, Frontal_Inf_Orb_R, Frontal_Inf_Tri_R |
| **d) Post-Instruction FCI > Control: Female > Male** | | | | | | | |
| 1 | R | 12 | -68 | 30 | 4104 | 3.729 | White Matter, Precuneus_R, Cuneus_R, Calcarine_R |
| **e) Pre-Instruction PK > Control: Male > Female** | | | | | | | |
| 1 | B | 6 | -68 | -42 | 7984 | 3.481 | |
| 2 | L | -46 | 10 | 28 | 4096 | 3.587 | White Matter, Frontal_Inf_Oper_L, Precentral_L, Frontal_Inf_Tri_L, Frontal_Mid_L, Rolandic_Oper_L |
| 3 | L | -26 | -66 | 46 | 3480 | 3.373 | White Matter, Parietal_Sup_L, Parietal_Inf_L, Occipital_Mid_L, Occipital_Sup_L |
| 4 | R | 32 | -88 | -2 | 2856 | 3.393 | White Matter, Occipital_Mid_R, Occipital_Inf_R, Lingual_R, Calcarine_R, Fusiform_R |
| 5 | L | -48 | -46 | 46 | 2608 | 3.388 | Parietal_Inf_L, Angular_L, White Matter |
| 6 | L | -48 | -56 | -12 | 1872 | 3.484 | White Matter, Temporal_Inf_L, Occipital_Inf_L, Fusiform_L, Temporal_Mid_L |
| **f) Pre-Instruction PK > Control: Female > Male** | | | | | | | |
| 1 | B | 0 | -84 | 12 | 6224 | 3.616 | Calcarine_L, Cuneus_R, Calcarine_R, Cuneus_L, Occipital_Sup_L, Lingual_L, White Matter |
| **g) Post-Instruction PK > Control: Male > Female** | | | | | | | |
| 1 | L | -50 | -50 | 48 | 5616 | 3.607 | Parietal_Inf_L, Parietal_Sup_L, Angular_L, SupraMarginal_L, White Matter |
| 2 | L | -54 | 14 | 18 | 1296 | 3.495 | White Matter, Frontal_Inf_Oper_L, Frontal_Inf_Tri_L, Precentral_L |
| **h) Post-Instruction PK > Control: Female > Male** | | | | | | | |
| 1 | B | 2 | -82 | 14 | 8296 | 3.535 | Calcarine_L, Calcarine_R, Cuneus_R, Cuneus_L, Lingual_L, Occipital_Sup_L, Occipital_Sup_R, Lingual_R, White Matter |
| 2 | L | -48 | 12 | -34 | 5416 | 3.648 | Temporal_Pole_Mid_L, Temporal_Mid_L, Temporal_Pole_Sup_L, Temporal_Inf_L, Frontal_Inf_Orb_L, Insula_L, Amygdala_L, White Matter |
| 3 | L | -12 | -60 | -8 | 3680 | 3.452 | Lingual_L, Cerebelum_4_5_L, Cerebelum_6_L, Calcarine_L, |

| | | | | | | | |
|---|---|---|---|---|---|---|---|
| | | | | | | | Precuneus_L, Vermis_4_5, White Matter |
| 4 | R | 14 | -56 | -12 | 3008 | 3.452 | White Matter, Cerebelum_4_5_R, Lingual_R, Cerebelum_6_R, Fusiform_R, Vermis_4_5 |
| 5 | L | -32 | -34 | -28 | 2096 | 3.614 | Fusiform_L, Cerebelum_4_5_L, Cerebelum_6_L, Cerebelum_Crus1_L, ParaHippocampal_L, White Matter |
| 6 | R | 40 | 20 | -40 | 1696 | 3.724 | Temporal_Pole_Mid_R, Temporal_Pole_Sup_R, Fusiform_R, White Matter |
| 7 | R | 58 | 2 | -32 | 1288 | 3.501 | Temporal_Mid_R, Temporal_Inf_R, Temporal_Pole_Mid_R, White Matter |

**Supplemental Table 6.** Brain Activation Results for Sex, Time, and Pedagogy Interactions. *Center of mass coordinates of significant regions of brain activation within the PK > Control contrast showing a-d) two-way (sex x time) interactions and e-f) three-way (pedagogy x sex x time) interactions. Cluster region labels are based off those reported by the IBASPM116 Human Brain Atlas. Coordinates are reported in MNI space.*

| Cluster | Hemisphere | Center of Mass (MNI space) | | | Cluster Extent ($mm^3$) | Mean Z Score | Labels |
|---|---|---|---|---|---|---|---|
| **a) PK > Control: Sex x Time Interactions** | | | | | | | |
| | | X | Y | Z | | | |
| 1 | B | 6 | -56 | -32 | 4912 | 3.368 | |
| 2 | L | -16 | -6 | 62 | 1608 | 3.555 | White Matter, Supp_Motor_Area_L, Frontal_Sup_L, Precentral_L, Paracentral_Lobule_L |
| 3 | L | -30 | 24 | 10 | 1488 | 3.443 | White Matter, Insula_L, Frontal_Inf_Tri_L, Frontal_Mid_L |
| 4 | R | 20 | -14 | 2 | 1368 | 3.409 | White Matter, Thalamus_R, Caudate_R, Pallidum_R, Putamen_R, Cerebro-Spinal Fluid |
| **b) PK > Control: Sex x Class x Time Interactions** | | | | | | | |
| 1 | L | -18 | -60 | 0 | 1856 | 3.347 | White Matter, Lingual_L, Calcarine_L, Precuneus_L, Cerebelum_4_5_L, Cerebro-Spinal Fluid |
| 2 | R | 14 | -82 | -52 | 1696 | 3.465 | |
| 3 | L | -40 | -56 | -20 | 1640 | 3.315 | White Matter, Fusiform_L, Cerebelum_6_L, Cerebelum_Crus1_L, Temporal_Inf_L, Occipital_Inf_L |
